# F-BAR Cdc15 Promotes Gef1-mediated Cdc42 Activation During Cytokinesis and Cell Polarization in *S. pombe*

**DOI:** 10.1101/552927

**Authors:** Brian S. Hercyk, Maitreyi E. Das

**Author notes:** Corresponding Author: Maitreyi Das, University of Tennessee, Knoxville, 415, Ken and Blaire Mossman Building, Knoxville, TN, 37996.

## Abstract

Cdc42, a Rho-family GTPase, is a master regulator of cell polarity. Recently it has been shown that Cdc42 also facilitates proper cytokinesis in the fission yeast, *Schizosaccharomyces pombe*. Cdc42 is activated by two partially redundant GEFs Gef1 and Scd1. Although both the GEFs activate Cdc42, their deletion mutants display distinct phenotypes, indicating that they are differentially regulated, by an unknown mechanism. During cytokinesis, Gef1 localizes to the division site and activates Cdc42 to initiate ring constriction and septum ingression. Here we report that the F-BAR domain containing Cdc15 promotes Gef1 localization to its functional sites. We show that *cdc15* promotes Gef1 association with the cytokinetic nodes to activate Cdc42 during ring assembly. Moreover, *cdc15* phospho-mutants phenocopy polarity phenotypes of *gef1* mutants. In a hypermorphic *cdc15* mutant, Gef1 localizes precociously to the division site, and is readily detected at the cortical patches and the cell cortex. Correspondingly, the hypermorphic *cdc15* mutant shows increased bipolarity during interphase and precocious Cdc42 activation at the division site during cytokinesis. Finally, loss of *gef1* in hypermorphic *cdc15* mutants abrogates the increased bipolarity and precocious Cdc42 activation phenotype. We did not see any change in the localization of the other GEF Scd1 in a Cdc15-dependent manner. Taken together our data indicates that Cdc15 promotes Cdc42 activation specifically via Gef1 localization to the division site to facilitate proper cytokinesis and to the cell cortex to promote bipolarity.

## INTRODUCTION

The conserved Cdc42 is a master regulator of polarized cell growth in fission yeast (Miller and Johnson 1994; Johnson 1999; Estravis *et al*. 2012; Das and Verde 2013). Recently, it has also been shown that Cdc42 has a role in cytokinesis, the final step in cell division (Wei *et al*. 2016). Through the regulation of actin and membrane trafficking, Cdc42 controls cellular processes such as growth, cell polarity, and cytokinesis (Martin *et al*. 2007; Harris and Tepass 2010; Estravis *et al*. 2011; Estravis *et al*. 2012). Given the complexities of these cellular processes, Cdc42 activation needs to be precisely regulated in a spatiotemporal manner. A prime example of this precise regulation is the oscillation of Cdc42 activation between the two cells ends during bipolar growth (Das *et al*. 2012; Das and Verde 2013). Disrupting Cdc42 activation patterns leads to defects in cell shape and cytokinesis (Das *et al*. 2012; Wei *et al*. 2016; Onwubiko *et al*. 2019). While much is known about how Cdc42 promotes actin organization and polarization, the spatiotemporal manner in which regulation of Cdc42 is fine-tuned is not well understood.

Cdc42 is activated by GEFs (guanine nucleotide exchange factors) which exchange GDP for GTP, and inactivated by GAPs (GTPase activating proteins) which enhance the intrinsic rate of GTP hydrolysis (Bos *et al*. 2007). Fission yeasts have two GEFs, Scd1 and Gef1 that control polarization and cytokinesis (Chang *et al*. 1994; Coll *et al*. 2003). While both the GEFs activate Cdc42 and their double deletion is not viable (Coll *et al*. 2003; Hirota *et al*. 2003), *scd1Δ* and *gef1Δ* mutants exhibit distinct phenotypes, indicating that they differentially activate Cdc42. *scd1Δ* cells are depolarized and exhibit defects in septum morphology (Chang *et al*. 1994; Wei *et al*. 2016). In contrast, *gef1Δ* mutants exhibit monopolar growth and a delayed onset of ring constriction (Coll *et al*. 2003; Das *et al*. 2015; Wei *et al*. 2016; Onwubiko *et al*. 2019). This suggests that the two GEFs allow for distinct Cdc42 activation patterns, which regulate different aspects of cell polarity establishment and cytokinesis. It is unclear how the two Cdc42 GEFs result in distinct phenotypes given they both activate the same GTPase. One potential explanation could be differential regulation of these GEFs. Indeed, during cytokinesis, first Gef1 is recruited to the membrane proximal to the actomyosin ring where it activates Cdc42 to promote timely onset of ring constriction and septum initiation (Wei *et al*. 2016). Next, Scd1 is recruited to the ingressing membrane to promote proper septum maturation (Wei *et al*. 2016).

It is unknown what gives rise to the temporal recruitment pattern of the GEFs. How does Gef1, but not Scd1, initially localize to the division site to activate Cdc42 in a timely manner? Gef1 contains an N-BAR domain that is required for its activity but not for its localization (Das *et al*. 2015). The N-terminal region of Gef1 is necessary and sufficient for its localization (Das *et al*. 2015). Phosphorylation of the N-terminal region by Orb6 kinase generates a 14-3-3 binding site that results in the sequestration of Gef1 in the cytoplasm (Das *et al*. 2009; Das *et al*. 2015). While it is known how Gef1 is removed from its site of action, it is unclear what recruits Gef1 to these sites.

Here we show that Gef1 localization to its site of action is aided by the F-BAR protein Cdc15. Cdc15 localizes to endocytic patches during interphase and to the division site, where it scaffolds the actomyosin ring (Wu *et al*. 2003; Arasada and Pollard 2011; McDonald *et al*. 2017). We report that Gef1 localizes to cortical nodes at the division site during ring assembly in a cdc15-dependent manner. Similarly, we find that *cdc15* promotes Gef1 localization to the cortical patches and cell tips. We show that *cdc15* phospho-mutants phenocopy *gef1* polarity phenotypes. A hypermorphic *cdc15* allele shows precocious Cdc42 activation at the division site during cytokinesis and increased bipolarity during interphase. Finally, we show that enhanced bipolarity and premature Cdc42 activation is abrogated upon deletion of *gef1* in the hypermorphic *cdc15* mutant. This indicates that *cdc15* promotes Cdc42 activation through *gef1*. We did not see any change in the localization of the other GEF Scd1 in a cdc15-dependent manner. Taken together our data indicates that Cdc15 specifically promotes Gef1 localization to the division site and the cell cortex to promote Cdc42 activation.

## MATERIALS AND METHODS

### Strains and cell culture

The S. *pombe* strains used in this study are listed in Supplemental Table S1. All strains are isogenic to the original strain PN567. Cells were cultured in yeast extract (YE) medium and grown exponentially at 25°C, unless specified otherwise. Standard techniques were used for genetic manipulation and analysis (Moreno *et al*. 1991). Cells were grown exponentially for at least 3 rounds of eight generations before imaging.

### Microscopy

Cells were imaged at room temperature (23–25°C) with an Olympus IX83 microscope equipped with a VTHawk two-dimensional array laser scanning confocal microscopy system (Visitech International, Sunderland, UK), electron-multiplying charge-coupled device digital camera (Hamamatsu, Hamamatsu City, Japan), and 100×/numerical aperture 1.49 UAPO lens (Olympus, Tokyo, Japan). Images were acquired with MetaMorph (Molecular Devices, Sunnyvale, CA) and analyzed by ImageJ (National Institutes of Health, Bethesda, MD (Schneider *et al*. 2012)). For still and z-series imaging, the cells were mounted directly on glass slides with a #1.5 coverslip (Fisher Scientific, Waltham, MA) and imaged immediately; fresh slides were prepared every 10 minutes. Z-series images were acquired with a depth interval of 0.4 μm. For time-lapse images, the cells were placed in 3.5-mm glass-bottom culture dishes (MatTek, Ashland, MA) and overlaid with YE medium plus 0.6% agar with 100μM ascorbic acid as an antioxidant to minimize toxicity to the cell, as reported previously (Frigault *et al*. 2009; Wei *et al*. 2017).

### Analysis of fluorescent intensity

Mutants expressing fluorescent proteins were harvested from mid-log phase cultures at OD_(595)_ 0.5 and imaged on slides. Depending on the mutant and the fluorophore, 16-18 z-planes were collected at a z-interval of 0.4μm for either or both the 488nm and 561nm channels. The respective controls were grown and imaged in an identical manner. ImageJ was used to generate sum projections from the z-series, and to measure the fluorescence intensity of a selected region. The cytoplasmic fluorescence of the same cell was subtracted to generate the normalized intensity. Mean normalized intensity was calculated for each image from all measurable cells (n>5) within each field.

### Statistical tests

Statistical tests were performed using GraphPad Prism software. When comparing two samples, a student’s t-test (two-tailed, unequal variance) was used to determine significance. When comparing three or more samples, one-way ANOVA was used, followed by a Tukey’s multiple comparisons post-hoc test to determine individual p-values.

### Cell staining

To stain the septum and cell wall, cells were stained in YE liquid with 50 μg/ml Calcofluor White M2R (Sigma-Aldrich, St. Louis, MO) at room temperature.

### Latrunculin A treatment

Cells were treated with 10 μM latrunculin A in dimethyl sulfoxide (DMSO) in YE medium for 30 min before imaging. Control cells were treated with only 0.1% DMSO in YE medium.

### Analysis of *sin* and *cdc12* mutants

*plo1-25, sid2-250,* and control cells were grown in YE at 25°C to OD 0.2, then shifted to the restrictive temperature at 35.5°C. Slides were then prepared and imaged from the cultures at 0, 1, 2, and 4 hour time points. Cells expressing cdc12ΔC-GFP were initially grown in EMM (Edinburgh minimal medium) with 150μM Thiamine. Induction of cdc12ΔC-GFP expression was performed as described previously (Yonetani and Chang 2010). Briefly, cultures were harvested by low speed centrifugation, rinsed, and then grown in EMM without thiamine for 18 hours prior to imaging.

## RESULTS

### Gef1 localizes to cytokinetic nodes

While we have previously characterized the distinct localization pattern and phenotypes of the Cdc42 GEFs, Gef1 and Scd1 during cytokinesis (Wei *et al*. 2016), how they are recruited to the division site at the appropriate time is unknown. Since Gef1 is detectable at the membrane proximal to the assembled actomyosin ring, we posited that the ring is required for Gef1 localization. To test this, we treated cells with 10μM Latrunculin A (LatA) for 30 min to depolymerize actin structures, then observed the localization of Gef1-mNeonGreen (Gef1-mNG). Gef1-mNG localizes to the membrane proximal to the actomyosin ring, marked by Rlc1-tdTomato, in mock DMSO treated cells (Fig. 1A). Rlc1-tdTomato rings fragment upon treatment with LatA, as does Gef1-mNG, indicating that an intact ring is necessary for proper Gef1 localization. We observe that upon LatA treatment, Gef1-mNG does not diffuse away into the cytosol, but instead localizes to cortical nodes about the cortex with Rlc1-tdTomato. Upon closer examination of these nodes, one population of Gef1 can be seen to partially colocalize with Rlc1, while the other population of Gef1 puncta do not overlap with Rlc1 (Fig. B). These findings indicate that Gef1 may interact with or be recruited by one of the proteins within these cortical nodes.

**Figure 1.**
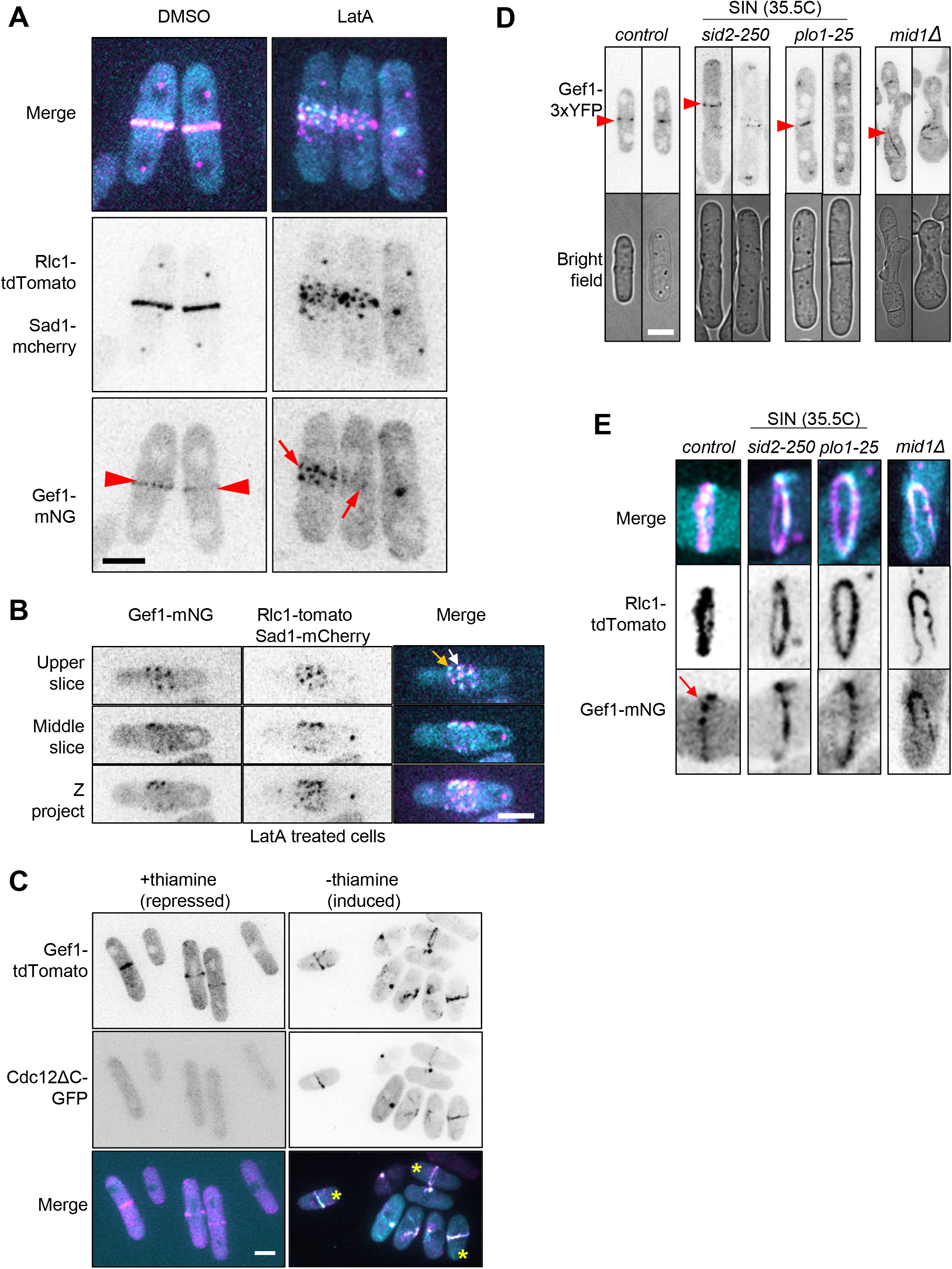
Gef1 associates with cytokinetic nodes. **(A)** Gef1-mNG and Rlc1-tdTomato localization was examined in cells treated with 10μM LatA for 30 min. In DMSO treated control cells, Gef1 (red arrowheads) localizes normally to the actomyosin ring. Upon treatment with the actin depolymerizing drug LatA, the ring fragments and Gef1 appears to localize to cortical nodes at division site (red arrows). **(B)** Top and middle z-series show node like organization of Gef1-mNG and Rlc1-tdTomato about the cortex at the division site in cells treated with LatA. White arrow shows colocalized Gef1 and Rlc1, while orange arrow shows Gef1 alone. **(C)** Expression of cdc12ΔC-GFP induces ectopic actomyosin ring formation and constriction in interphase cells. Gef1-tdTomato ectopically localizes to these rings that form in interphase (yellow asterisks). **(D)** Gef1 localizes to aberrant ring-like structures formed in *sin* and *mid1Δ* mutants. Indicated genotypes were shifted to the restrictive temperature of 35.5°C for 4 hours. Top row: Inverted max projections of Gef1-3xYFP (red arrowheads). Bottom row: Brightfield images of the representative images above. **(E)** Gef1 colocalizes with Rlc1-tdTomato in the aberrant rings formed in *sin* and *mid1Δ* mutants. Red arrow shows node like organization of Gef1 at the actomyosin ring in *sid2*^+^ *plo1*^+^ *mid1*^+^ control cells. Merge of the division site of *control* and *sin* and *mid1Δ* mutant cells expressing Gef1-mNG and Rlc1-tdTomato. Scale bars represent 5μm.

Since Gef1 promotes timely onset of ring constriction (Wei *et al*. 2016), we asked if Gef1 localization itself was under a temporal control during cytokinesis. Given that Gef1 arrives at the division site during anaphase as the actomyosin ring assembles (Wei *et al*. 2016), we asked whether Gef1 is recruited in a cell cycle-dependent manner. To test this, we induced ectopic ring formation in interphase cells using the constitutively active formin mutant, cdc12ΔC-GFP (Yonetani and Chang 2010). In the presence of thiamine, cdc12ΔC-GFP expression is repressed. In these conditions, Gef1-tdTomato localizes to the division site of mitotic cells, which are approximately 14μm in length (Fig. 1C). However, induction of cdc12ΔC-GFP expression results in the formation of ectopic rings in mono-nucleate interphase cells less than 10μm long, to which Gef1-tdTomato localizes (Fig. 1C). This indicates that Gef1 localization to the ring is not cell cycle-dependent, but rather that formation of the actomyosin ring is sufficient for Gef1 localization.

Next, we asked what pathway recruits Gef1 to the division site. The Septation Initiation Network (SIN) is a protein signaling network that coordinates the timing of cytokinesis with chromosome segregation (Roberts-Galbraith and Gould 2008; Johnson *et al*. 2012; Simanis 2015). The SIN pathway recruits and activates proteins involved in ring constriction and the coordinated process of septum formation (Jin *et al*. 2006; Roberts-Galbraith *et al*. 2010; Bohnert *et al*. 2013). To determine whether the SIN is required for Gef1 localization to the division site, we examined the localization of Gef1-3YFP in two SIN protein kinase *ts* mutants, *plo1-25 and sid2-250* (Bahler *et al*. 1998; Jin *et al*. 2006; Hachet and Simanis 2008). In *plo1-25 and sid2-250,* Gef1-3xYFP localizes normally to the division site at the permissive temperature of 25°C. Surprisingly, Gef1 - 3YFP still localizes to ring like structures in *plo1-25 and sid2-250* cells shifted to the restrictive temperature of 35.5°C for 1, 2, or 4 hours (Fig. 1D). We imaged *plo1-25 and sid2-250* cells expressing Gef1-mNG and the ring marker Rlc1-tdTomato to better visualize the ring-like Gef1 structures and to determine whether these structures represented components of the actomyosin ring. Indeed, Gef1-mNG colocalizes with Rlc1-tdTomato in cells shifted to 35.5°C for 1, 2, or 4 hours, demonstrating that Gef1 recruitment to the actomyosin ring in not dependent upon the SIN pathway (Fig. 1E). Since we had observed Gef1 localization to cortical nodes, we examined whether Gef1 recruitment was Mid1-dependent. Mid1 is an anillin-like protein that is exported from the nucleus to form cortical nodes that define the division plane (Bahler *et al*. 1998; Paoletti and Chang 2000). It is to these nodes that various contractile ring components are recruited, before coalescing to form the actomyosin ring (Coffman *et al*. 2009; Laporte *et al*. 2011). In *mid1Δ* cells, Gef1-3xYFP localizes to misplaced, extended ringlike structures (Fig.1D). Gef1-mNG and Rlc1-tdTomato colocalize at these extended ring-like structures, similarly to the *sin* mutants (Fig.1E). This demonstrates that the early node protein Mid1 is not required for the localization of Gef1 to the actomyosin ring.

Closer examination of the division site prior to ring constriction shows patch like Gef1-mNG distribution rather than a continuous ring (Fig. 1A and E). This is in agreement with our observation that Gef1 associates with cytokinetic nodes at the division site upon LatA treatment.

### Gef1-dependent Cdc42 activation appears at the division site prior to ring assembly

The observation that Gef1 appears to localize to node like patches at the division site upon LatA treatment suggests that Gef1 may also be present in nodes during ring assembly. However, Gef1 is localized mainly in the cytoplasm and cannot be easily detected when present in small quantities at the membrane. Gef1 is the first GEF to localize to the division site and activate Cdc42 (Wei *et al*. 2016). To determine if Gef1 indeed localizes to the division site prior to ring assembly, we carefully examined Gef1-mediated Cdc42 activity during ring assembly. We monitored Cdc42 activity with the CRIB-3xGFP bio-probe that specifically binds to active GTP-bound Cdc42 (Tatebe *et al*. 2008). In *gef1*^+^ cells, CRIB-3xGFP first appears as a broad band at the division site, as it is lost from the nucleus, 8 minutes after the Sad1-mCherry labelled spindle pole bodies (SBP) separates (Fig. 2A, red arrowhead, 2D). However, the actomyosin ring, visualized by Rlc1-tdTtomato, does not fully assemble for another 4 minutes (Fig.2A, blue arrowhead). This suggests that Cdc42 is activated at the cortical nodes before they completely condense to form the cytokinetic ring. In contrast, CRIB-3xGFP does not become active at the division site until ~44 minutes after SPB separation (Fig. 2B, red arrowhead, 2D). Thus, although Gef1 cannot be directly detected at the division site during this period, our findings suggest that Gef1 specifically activates Cdc42 as the ring assembles (Fig. 2B).

**Figure 2.**
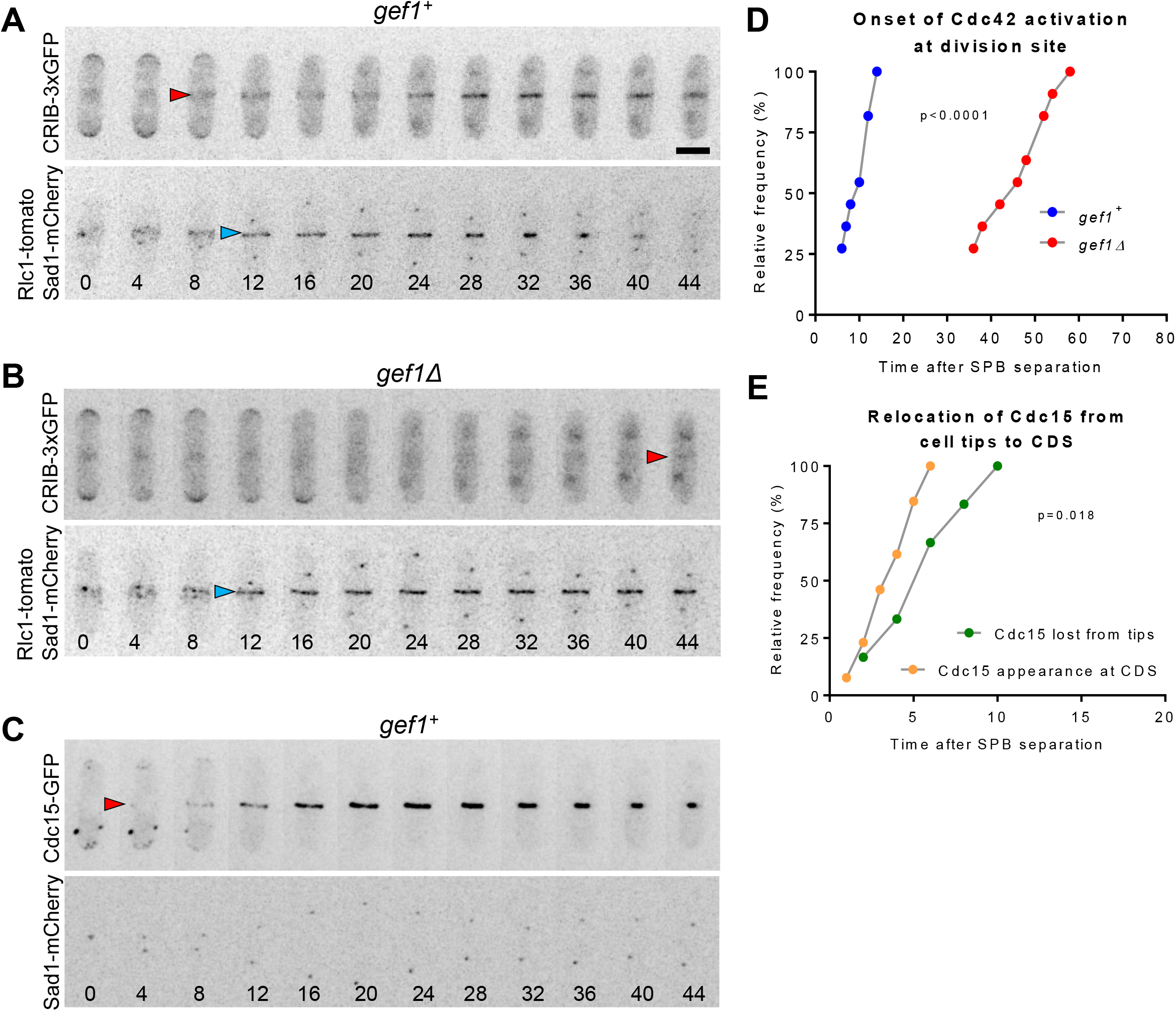
Cdc42 activation at the division site initiates during actomyosin ring formation. **(A)** In *gef1*^+^ cells, CRIB-3xGFP appears at the division site prior to ring assembly. **(B)** In *gef1Δ* cells, CRIB-3xGFP does not appear at the division site until the onset of ring constriction. **(C)** Cdc15-GFP appears at the division site and begins to condense into the ring just prior to Cdc42 activation. Montages are inverted z-projections of the same cells imaged over time. Numbers beneath montages represent time in minutes with respect to SPB separation. Red arrowheads mark the time at which CRIB-3xGFP is first detected at the division site (A and B) or Cdc15-GFP (C). Blue arrowheads mark ring formation. **(D)** Frequency distribution plot of the percentage of cells in the indicated strains with CRIB-3xGFP at the division site as a function of time since SPB separation. **(E)** Frequency distribution plot of the relocation of Cdc15 from the tips to the division site as a function of time since SPB separation. Reported p-values from Student’s t-test. Scale bar represents 5μm.

While our data suggests that Gef1 associates with cortical nodes, its localization is independent of the early node protein, Mid1. Since Gef1-mediated Cdc42 activation initiates during ring formation, we posit that a later node protein may recruit Gef1 to the division site. One of the last proteins recruited to the cortical nodes before ring formation is the F-BAR Cdc15 (Wu *et al*. 2003). During cytokinesis Cdc15 is redistributed from the cell tips to the division site. We find that while Cdc15 localizes to the division site ~ 4 minutes after SPB separation, patches of Cdc15 remain at the polarized growth regions until ~10 minutes after SPB separation (Fig. 2C, 2D). This suggests that Cdc42 is activated at the division site as Cdc15 is redistributed within the cell. Since Cdc42 activation is solely Gef1-mediated during this period, we asked whether Cdc15 may recruit Gef1 to the division site to activate Cdc42. A recent report indicates that Gef1 regulates Cdc15 distribution along the actomyosin ring (Onwubiko *et al*. 2019). It is possible that Gef1 via a feedback mechanism depends on Cdc15 for its localization.

### Cdc15 promotes Gef1 localization to the division site

Cdc15 associates with the membrane via its F-BAR domain and acts as a scaffold that associates with proteins at the actomyosin ring (McDonald *et al*. 2015; Ren *et al*. 2015; McDonald *et al*. 2017). The scaffolding ability of Cdc15 is primarily conferred through its C-terminal SH3 domain, through which it interacts with other proteins (Roberts-Galbraith *et al*. 2009; Ren *et al*. 2015). While *cdc15* is essential for fission yeast, a *cdc15ΔSH3* mutant is viable but displays defects in septum ingression and ring constriction (Roberts-Galbraith *et al*. 2009). Similar to *gef1Δ* mutants, onset of ring constriction and Bgs1 localization to the division site is delayed in *cdc15ΔSH3* mutants (Roberts-Galbraith *et al*. 2009; Arasada and Pollard 2014; Cortes *et al*. 2015; Wei *et al*. 2016). In these mutants the Cdc15 interacting proteins are partially lost from the division site (Ren *et al*. 2015). We posit that Cdc15 may promote Gef1 localization to the division site through interaction with its SH3 domain. To test this, we examined Gef1-tdTomato localization to rings that were assembled, but not constricting, in cells expressing either Cdc15-GFP or cdc15ΔSH3-GFP. Gef1-tdTomato is present in ~70% of Cdc15-GFP rings, while Gef1-tdTomato is present in only ~40% of cdc15ΔSH3-GFP rings (Fig. 3A and Fig. 3B). Furthermore, Gef1-tdTomato fluorescent intensity is also reduced at the assembled rings of the *cdc15ΔSH3* mutant, with a relative intensity of only 76% that of *cdc15*^+^ cells (Fig. 3A, 3C, Table 1). We find that Gef1-tdTomato localizes to the division site in cells with a minimum SPB distance of 3μm in *cdc15*^+^ cells. In contrast, in *cdc15ΔSH3* mutants Gef1-tdTomato appears at the division site with a minimum SPB distance of 7μm (Table 1). Moreover, in *cdc15*^+^ cells 61% of cells in anaphase B displayed Gef1-tdTomato at the division site, while in *cdc15ΔSH3* mutants only 12% of anaphase B cells showed Gef1 localization (Table 1). While Gef1 localization to assembled rings is initially impaired in cells expressing cdc15ΔSH3-GFP, all constricting rings have Gef1-tdTomoto (Table 1). This suggests that Cdc15 may either initially stabilize Gef1 or promote its localization to the division site.

**Figure 3.**
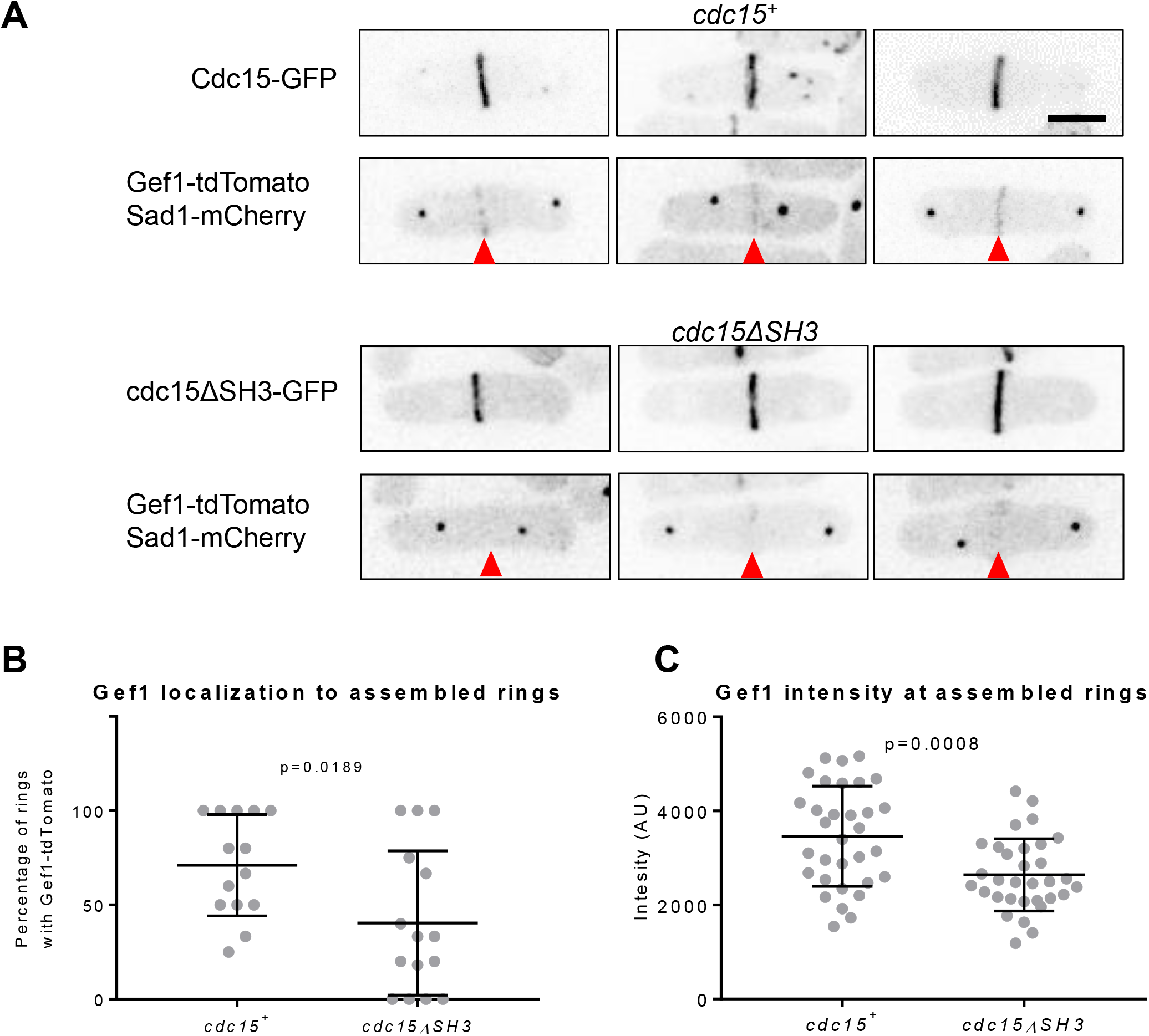
Cdc15 promotes Gef1 localization to the division site. **(A)** Inverted max projections of *cdc15*^+^ and *cdc15ΔSH3* expressing Cdc15-GFP, Gef1-tdTomato, and Sad1-mCherry. Red arrowheads mark the division site. **(B)** Quantification of fields of cells of the indicated genotypes that have Gef1-tdTomato present at the assembled actomyosin ring. **(C)** Quantification of Gef1-tdTomato intensity at assembled, but not constricting, rings in the indicated genotypes. Reported p-values from Student’s t-test. Scale bar represents 5μm.

**Table 1.** Characterization of Gef1 recruitment to the division site in a *cdc75*-dependent manner

|  | SPB distance<br>at which Gef1<br>first appears at<br>CDS | % of Anaphase<br>B cells with<br>Gef1 at CDS | % of cells with<br>assembled<br>rings with Gef1<br>at CDS | % of cells with<br>assembled<br>rings with Gef1<br>at CDS | Relative Gef1-tdTomato<br>intensity |
| --- | --- | --- | --- | --- | --- |
| *cdc15*<sup>+</sup> | 3μm<br>N=26 | 61%<br>N=26 | 78%<br>N=56 | 100%<br>N=24 | 1.0<br>N=32 |
| *cdc15ΔSH3* | 7μm<br>n=26 | 12%<br>n=26 | 48%<br>n=37 | 100%<br>n=43 | 0.76<br>n=32 |

### *cdc15* phenocopies *gef1* polarity phenotypes

Our data indicates a functional relationship between Gef1 and Cdc15 during cytokinesis. This is further supported by the fact that *cdc15*Δ*SH3* and *gef1* share a common phenotype: a delay in the onset of ring constriction and Bgs1 localization at the division site (Arasada and Pollard 2014; Cortes *et al*. 2015; Wei *et al*. 2016). It is possible that during cytokinesis Cdc15 recruits Bgs1 to the division site through Gef1. Since *gef1* has been shown to regulate cell polarity, we asked if *cdc15* functionally interacts with Gef1 during cell polarization. *gef1Δ* cells are primarily monopolar, growing only from the old end (Coll *et al*. 2003; Das *et al*. 2015). In contrast, the hypermorphic allele *gef1S112A* exhibits precocious new end growth, producing primarily bipolar cells (Das *et al*. 2015). We asked if this functional relationship between Gef1 and Cdc15 is specific to cytokinesis, or whether it is also observed during polarized growth. Indeed, as compared to control cells *cdc15*Δ*SH3* mutants show decreased bipolarity in interphase cells, similar to that observed in *gef1*Δ cells (Fig. 4A, 4B). Next, we asked if an increase in bipolarity was also observed in *cdc15* mutants with increased cortical localization. When oligomerized, the F-BAR domain enables Cdc15 to properly interact with the membrane (Roberts-Galbraith *et al*. 2010; McDonald *et al*. 2015). Cdc15 is a phospho-protein where hyper-phosphorylation disrupts proper oligomerization and at least in part impairs function (Roberts-Galbraith *et al*. 2010). In contrast, the de-phosphorylated form of Cdc15 shows increased oligomerization and increased localization at cortical patches (Roberts-Galbraith *et al*. 2010). We find that similar to *gef1*Δ and *cdc15*Δ*SH3* mutants, the phosphomimetic *cdc15-27D* allele exhibits decreased bipolarity (Fig. 4A, 4B). Further, the non-phoshorylatable *cdc15-27A* allele is primarily bipolar, similar to *gef1S112A* mutants (Fig.4A, 4B). These results demonstrate that, as during cytokinesis, *cdc15* functionally interacts with *gef1* also during cell polarization.

**Figure 4.**
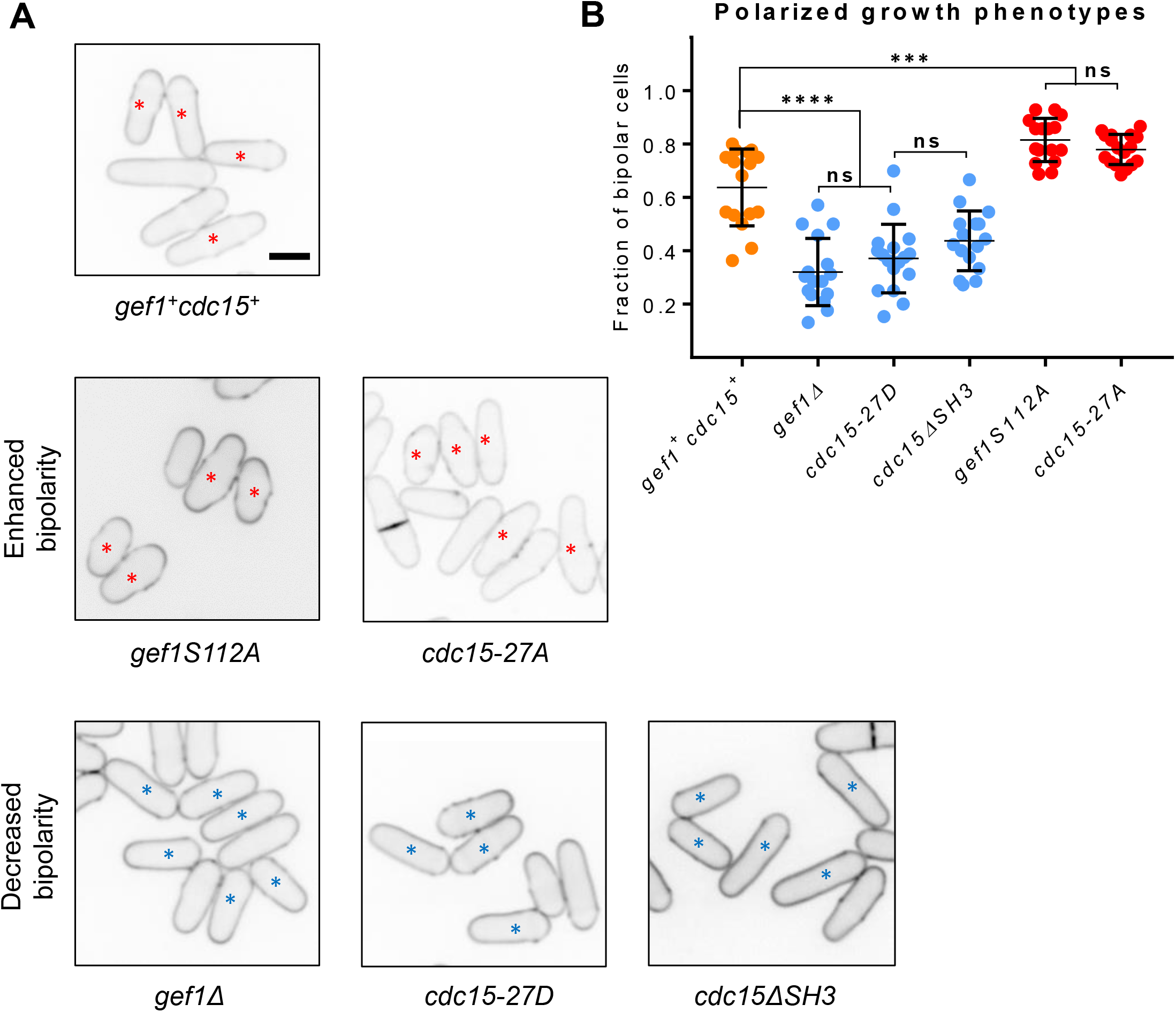
*cdc15* phenocopies *gef1* polarity phenotypes. **(A)** Representative images of the indicated genotypes stained with calcofluor to visualize polarized growth. Red asterisks denote bipolar cells, blue asterisks mark monopolar cells. **(B)** Quantification of the polarized growth phenotypes in the indicated genotypes. (****, p<0.0001, ***, p<0.001, ns, not significant, p-values reported from ANOVA with Tukey’s multiple comparisons post hoc test). Scale bar represents 5μm.

### *cdc15-27A* enhances Gef1 localization at cortical patches and division site

Gef1 is predominantly a cytosolic GEF, at least during interphase, and only transiently localizes to sites of polarized growth (Das *et al*. 2015). Given that Cdc15 promotes the recruitment of Gef1 to the division site, we asked whether it may also promote its localization to the sites of polarized growth, as suggested by the polarity phenotypes exhibited by *cdc15* mutants. While Cdc15-GFP is clearly visible at endocytic patches at the cell tips, Gef1-tdTomato is seldom observed (Fig. 5A, i). The non-phoshorylatable *cdc15-27A* mutants tagged to GFP show increased localization at cortical patches during interphase. Correspondingly, in cells expressing cdc15-27A-GFP, Gef1-tdTomato is readily observed at the cell cortex (Fig. 5A, ii, iii). Moreover, we also observed colocalization of Gef1-tdTomato and cdc15-27A-GFP at the cortical patches (Fig. 5A, v, red arrow). Interestingly, in addition to these colocalized patches, some regions of the cortex contain only Gef1-tdTomato or Cdc15-GFP (Fig. 5A, iv, green and orange arrowheads respectively). Next, we asked if Cdc15 also promoted Gef1 localization to the division site. We observe Gef1-mediated Cdc42 activation at the division site well before Gef1 itself is detectable (Fig. 1A). Similar to a previous report, Gef1-tdTomato can be detected at the division site only in rings that have completed assembly (Wei *et al*. 2016). We find that in *cdc15-27A* mutants, Gef1-tdTomato localizes to the division site before the ring completes assembly. Gef1-tdTomato colocalizes with cdc15-27A-GFP as the latter condenses into a ring, while it is not yet detectible at this stage in *cdc15*^+^ cells (Fig. 5B). Since it is hard to distinguish Gef1 signal at the division site from the cytoplasmic signal, it is not possible to precisely determine when Gef1 localizes to the division site by time lapse microscopy. However, we find that Gef1-tdTomato is detectable at cortical nodes in the medial region of interphase cells (Fig. 5C, yellow circles). Thus, it is possible that Gef1 localizes earlier to the division site in *cdc15-27A* mutants.

**Figure 5.**
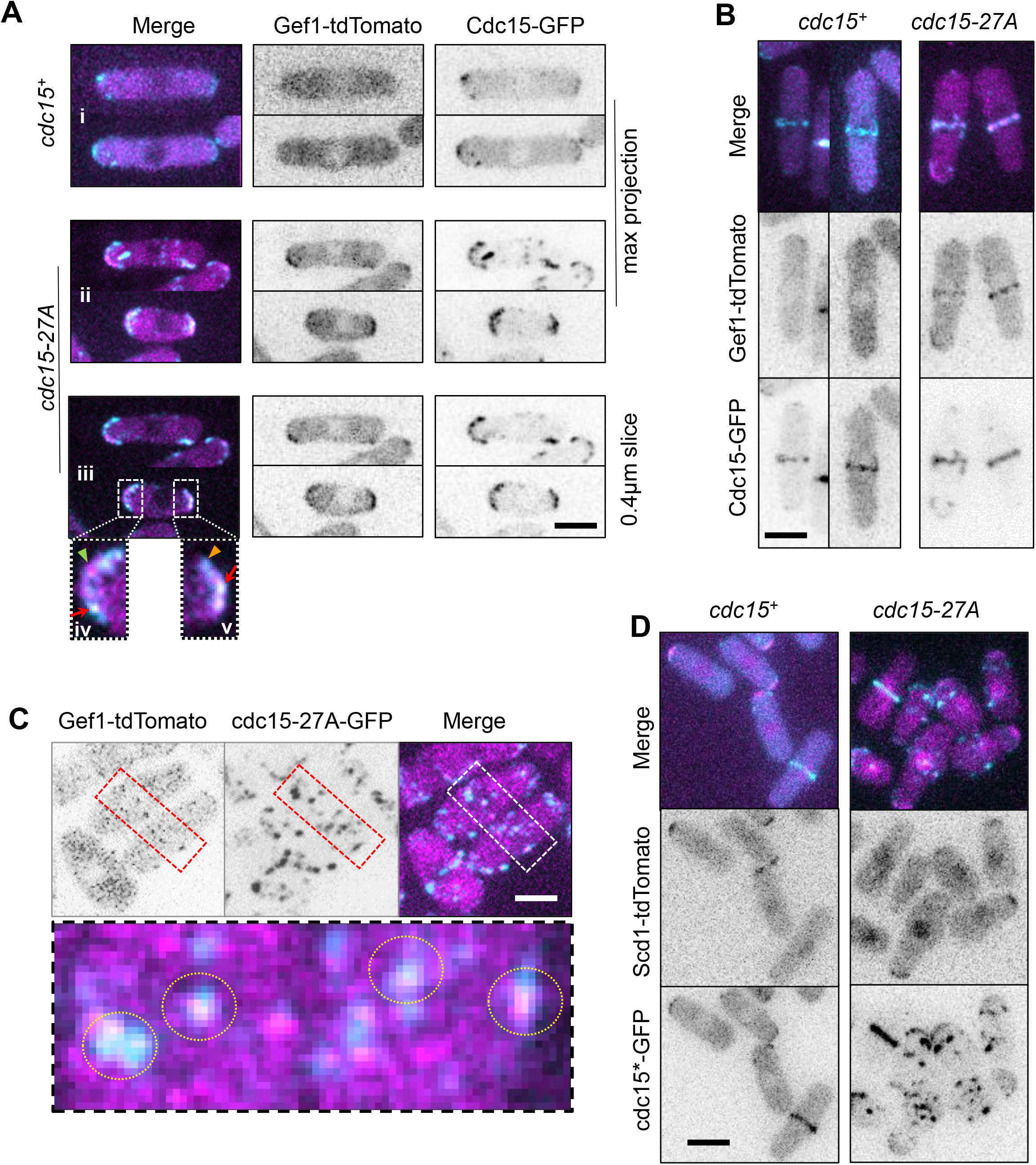
*cdc15-27A* enhances Gef1 localization at cortical patches. **(A)** Gef1-tdTomato and Cdc15-GFP localization to cortical patches in interphase *cdc15*^+^ and *cdc15-27A* cells. i and ii are max projections, while iii is a single 0.4μm z-plane of the same cell in ii. Insets iv and v are enlarged regions of the cell poles marked by white boxes. Red arrows indicate colocalization of Gef1 and Cdc15 patch. Green arrowhead indicates a Gef1 patch that does not colocalize with Cdc15. Orange arrowhead indicates a Cdc15 patch that does not colocalize with Gef1. **(B)** Gef1-tdTomato and Cdc15-GFP localization to the division site in *cdc15*^+^ and *cdc15-27A* cells. **(C)** Gef1-tdTomato and Cdc15-GFP localization to cortical patches at the division site in interphase *cdc15*^+^ and *cdc15-27A* cells. Yellow circles indicate colocalization of Gef1 and Cdc15 patch. Inset is an enlarged region of nascent division site marked by white box. **(D)** Scd1-tdTomato localization in cdc15-GFP and cdc15-27A-GFP expressing cells. Scale bar represents 5μm.

Next, we addressed whether Cdc15 specifically interacts with Gef1 or whether this interaction also extends to the other Cdc42 GEF. We examined the localization of the Cdc42 GEF Scd1 in *cdc15-27A* mutants. Under normal conditions, Scd1-tdTomato localization appears as a cap at the cell tips during interphase (Kelly and Nurse 2011; Das *et al*. 2012). We find that Scd1-tdTomoto localization in *cdc15-27A* cells does not differ from *cdc15*^+^ cells and do not localize to interphase cortical patches and nodes (Fig. 5D). This indicates that Cdc15 specifically promotes Gef1 localization during cytokinesis and cell polarization.

### Cdc15 promotes Gef1-mediated Cdc42 activation

Given that Gef1 precociously localizes to nodes at the division site of cells expressing *cdc15-27A*, we asked whether this was concomitant with precocious Cdc42 activation. Normally Cdc42 activity, visualized by CRIB-3xGFP, first appears at the division site only after the cell initiates anaphase A (Wei *et al*. 2016). However, we find that in *cdc15-27A* mutants, CRIB-3xGFP signal was visible at the cell medial region, even before the Sad1-mCherry labelled SPB separated. In these cells CRIB-3xGFP signal appeared as a broad band that overlapped with the nucleus (Fig. 6A). Next, we performed time lapse microscopy to determine when Cdc42 was activated at the division site in *cdc15-27A* mutants. Cdc42 is first activated ~10 minutes after SBP separation in *cdc15*^+^ cells. We find that in *cdc15-27A* mutants, Cdc42 is activated earlier at ~4 minutes after SPB separation, as determined by CRIB-3xGFP localization (Fig. 6B, red arrowhead, Supplementary Fig. S1). Further, similar to previous reports, in *cdc15*^+^ cells CRIB-3xGFP signal at the division site appears concurrent to loss of signal from the cell tips. We find that in *cdc15-27A* mutants, CRIB-3xGFP signal at the division site appears well before the signal is lost from the cell tips (Fig. 6B, yellow asterisk). While CRIB-3xGFP signal at the cell medial region in clearly detected in cells with a single SPB by still imaging, we did not detect CRIB-3xGFP signal at the division site prior to SPB separation by time lapse imaging. This could be due to low abundance or photobleaching of the signal, or it is possible that Cdc42 is only transiently activated at the medial region during interphase in *cdc15-27A* cells.

**Figure 6.**
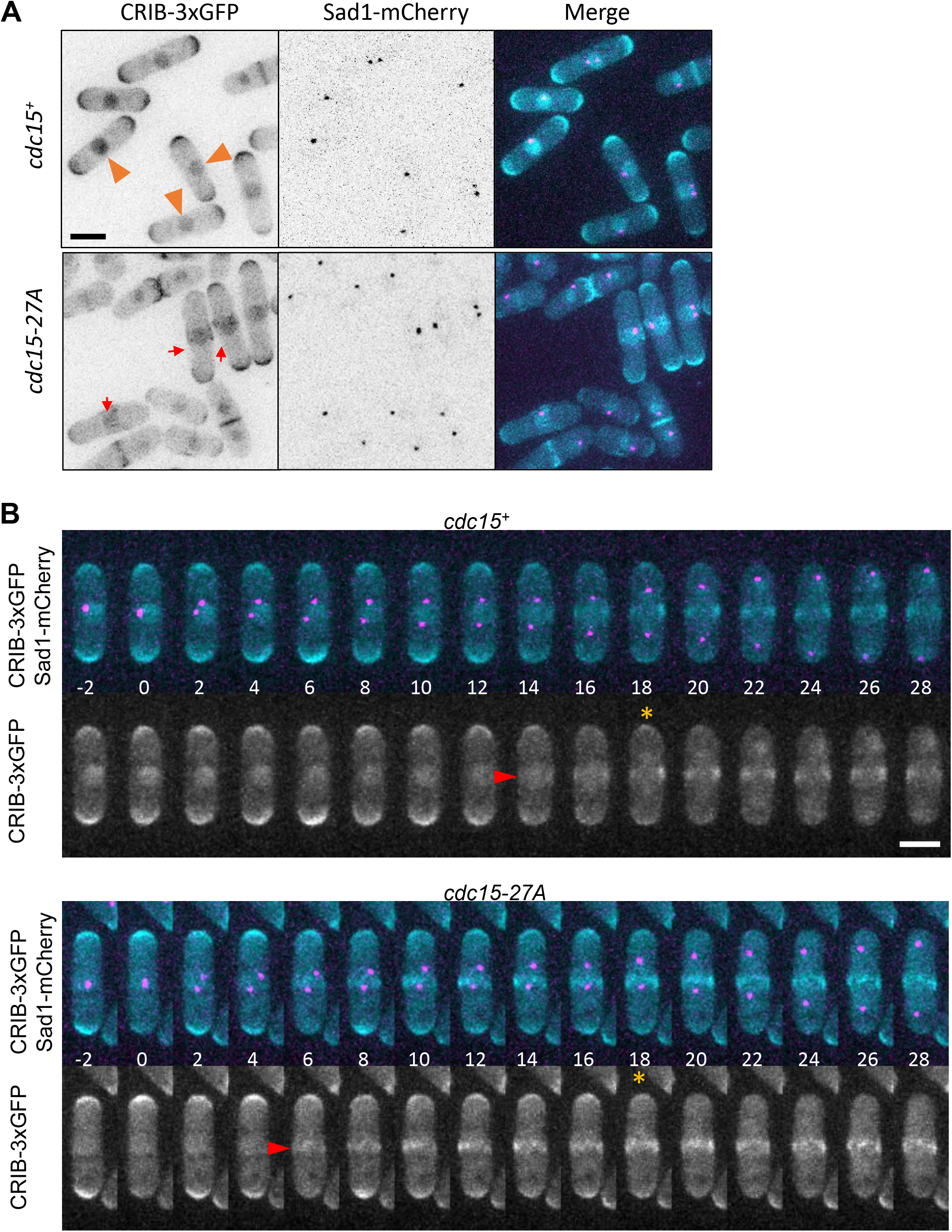
Cdc42 is prematurely activated in *cdc15-27A* cells during cytokinesis. **(A)** Inverted max projections of the indicated genotypes expressing CRIB-3xGFP and Sad1-mCherry. Orange arrowheads mark interphase cells without CRIB-3xGFP at the division site. Red arrows mark interphase cells with premature Cdc42 activation at the division site. **(B)** Time lapse montages of *cdc15*^+^ and *cdc15-27A* cells expressing CRIB-3xGFP and Sad1-mCherry. Red arrowheads mark onset of Cdc42 activation at the division site. Orange asterisks mark last time points before Cdc42 is completely lost from the cell tips. Numbers beneath montages represent time in minutes with respect to SPB separation. Scale bar represents 5μm.

Finally, we asked if the premature CRIB-3xGFP signal at the division site and the increased bipolarity observed in *cdc15-27A* mutants was due to Gef1-mediated Cdc42 activation. To test this, we deleted *gef1* in *cdc15-27A* mutants. While *cdc15-27A* mutants display an increased number of bipolar cells, the number of bipolar cells in the *gef1*Δ *cdc15-27A* double mutant was significantly reduced (p<0.0001) and similar to that observed in *gef1*Δ mutants (Fig. 7A, 7C). Furthermore, *cdc15-27A* cells display an increase in bipolar CRIB-3xGFP localization at the cell tips, relative to *cdc15*^+^ cells (Fig. 7B, 7D, p=0.039). This is consistent with our calcofluor data, indicating that bipolar growth is enhanced by *cdc15-27A*. Deletion of *gef1* in *cdc15-27A* mutants reduces bipolar CRIB-3xGFP localization, similar to that observed in *gef1*Δ cells (Fig. 7B, D, p<0.0001). Likewise, premature Cdc42 activation at the division site in *cdc15-27A* mutants is also abrogated in *gef1*Δ *cdc15-27A* cells. In *gef1Δ cdc15-27A* mutants CRIB-3xGFP did not appear at the division site until ~45minutes after SPB separation, as was also observed in *gef1Δ* (Fig. 7E; Supplementary Fig. S1). Together, these results indicate that Cdc15 promotes Cdc42 activation during cytokinesis and cell polarization via Gef1 localization.

**Figure 7.**
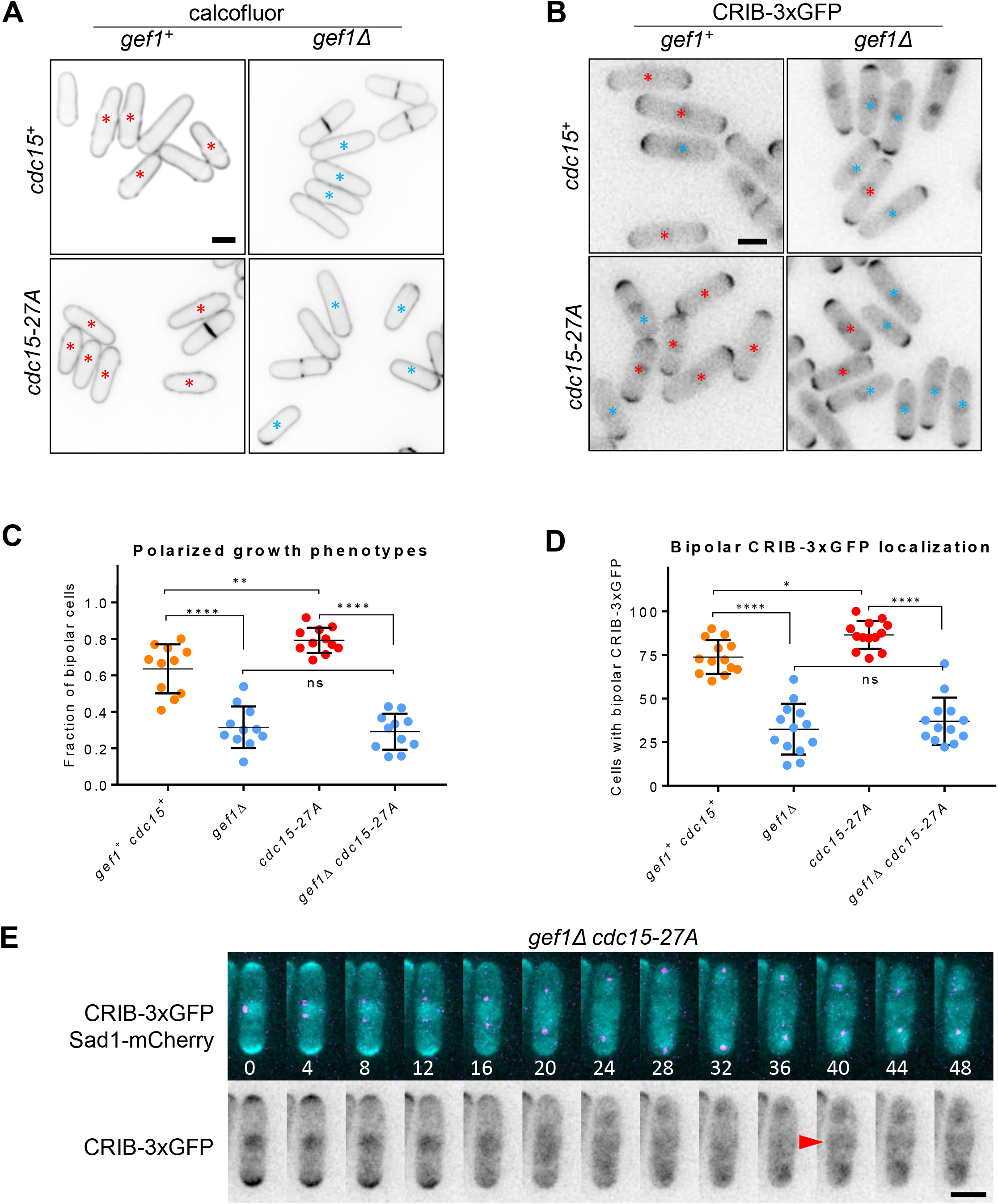
Cdc15 promotes Gef1-mediated Cdc42 activation. **(A)** Representative images of the indicated genotypes stained with calcofluor to visualize polarized growth. Red and blue asterisks denote bipolar and monopolar cells, respectively. **(B)** Quantification of the polarized growth phenotypes in the indicated genotypes. **(C)** Inverted max projections of the indicated strains expressing CRIB-3xGFP. Red asterisks mark cells with bipolar CRIB, blue asterisks show cells with monopolar CRIB localization. **(D)** Quantification of CRIB-3xGFP localization in the indicated genotypes. **(E)** Time lapse montages of *gef1Δ cdc15-27A* cells expressing CRIB-3xGFP and Sad1-mCherry. Red arrowheads mark onset of Cdc42 activation at the division site. Numbers beneath montages represent time in minutes with respect to SPB separation. (****, p<0.0001, ***, p<0.001, ns, not significant, one-way ANOVA with Tukey’s multiple comparisons post hoc test). Scale bar represents 5μm.

## DISCUSSION

The two Cdc42 GEFs, while partially redundant, show distinct phenotypes during cell polarity and cytokinesis (Chang *et al*. 1994; Coll *et al*. 2003; Wei *et al*. 2016). This suggests that the GEFs may be regulated in different ways to precisely activate Cdc42 at its site of function. Although the role of the Cdc42 GEF, Gef1 in cytokinesis and cell polarity is well established (Das and Verde 2013; Chiou *et al*. 2017), it is not clear how Gef1 localizes to its site of action. Here we show that Gef1, but not the other GEF Scd1, localizes to its site of action in a manner dependent on the F-BAR Cdc15.

Disintegration of the actomyosin ring by LatA treatment results in Gef1 localizing to the cytokinetic nodes suggesting that Gef1 associates with these structures. Gef1 does not become visible at the division site until the nodes coalesce into the actomyosin ring (Wei *et al*. 2016). However, Gef1-dependent Cdc42 activity can be observed a few minutes before the nodes fully coalesce to form the ring. Given that Gef1 is a low abundance protein, it is possible that Gef1 may be present at quantities beneath our detection limit at the cortical nodes during the initial stages of ring assembly.

Given the timing of Cdc42 activation, Gef1 appears to be recruited late during the ring assembly process. Thus, we looked at other proteins that are likewise recruited to the cytokinetic nodes during a similar time frame. It has previously been reported that the F-BAR protein Cdc15 is one of the last proteins to be recruited to the cytokinetic nodes before the ring assembles (Wu *et al*. 2003). We find that Cdc15 localizes to the division site shortly before Gef1-depdendent Cdc42 activity initiates. This seemed a likely candidate for Gef1 recruitment, as Cdc15 serves as a scaffold for many other proteins during cytokinesis. The Cdc15 scaffolding activity is dependent on its C-terminal SH3 domain (Ren *et al*. 2015). We asked whether Cdc15 recruited Gef1 to the division site via the SH3 domain. We find that Gef1 recruitment is delayed in *cdc15ΔSH3* cells, and Gef1 levels at the division site remain low throughout constriction. Thus, our data suggests that Gef1 localization to the division site is *cdc15* dependent. The F-BAR Imp2 also stabilizes proteins at the actomyosin ring via its SH3 domain. While loss of one of these SH3 domains is permissible, the actomyosin ring of *imp2ΔSH3 cdc15ΔSH3* double mutant does not retain its cohesion and disintegrates without constriction (Roberts-Galbraith *et al*. 2009; Ren *et al*. 2015). It is possible that like Cdc15, Imp2 may also recruit or stabilize Gef1 at the division site.

We have previously reported that the β-1,3-glucan synthase Bgs1, the septum synthesizing enzyme that drives membrane ingression, is delayed in *gef1Δ* cells (Wei *et al*. 2016). A similar defect is observed in *cdc15ΔSH3* (Roberts-Galbraith *et al*. 2009; Cortes *et al*. 2015). Given that Cdc15 promotes Gef1 localization to the division site, and that *cdc15ΔSH3* also exhibits the delayed onset of ring constriction (Roberts-Galbraith *et al*. 2009), characteristic of *gef1Δ* cells, we posited that Cdc15 acts upstream of Gef1 during cytokinesis. Apart from its role in cytokinesis, Gef1 is also required for proper cell polarity establishment (Coll *et al*. 2003). In fission yeast, immediately after division the cells grow in a monopolar manner from the old end and; as the cells reach a certain size, bipolarity ensues (Das *et al*. 2012; Das *et al*. 2015). Loss of *gef1* leads to a delay in initiation of bipolarity and as a result a large number of the cells in interphase are monopolar (Coll *et al*. 2003; Das *et al*. 2015). We asked if the relationship between Gef1 and Cdc15 was also conserved during cell polarity establishment. Thus, we examined the polarity phenotype of various *cdc15* mutants. Cdc15 is regulated via phosphorylation, and the phospho-mimetic allele *cdc15-27D* has been shown to adopt a closed conformation under cryo-EM, potentially reducing its ability to interact with other proteins (Roberts-Galbraith *et al*. 2010). Conversely, the non-phosphorylatable allele *cdc15-27A* adopts an open conformation that readily oligomerizes and is potentially hypermorphic (Roberts-Galbraith *et al*. 2010). While Gef1 mainly localizes to the cytoplasm, it cortical localization is enhanced in *gef1S112A* mutants rendering the cells bipolar. We find that the gain of function *cdc15-27A* mutant resembles *gef1S112A* mutants, in which the cells are predominantly bipolar. In contrast, *cdc15-27D* and *cdc15ΔSH3* mutants mimic *gef1Δ* mutants, in which cells are predominantly monopolar. Thus, *cdc15* phenocopies *gef1* during cytokinesis as well as in polarity establishment, providing further evidence of a functional interaction between these proteins.

A recent report suggests that Gef1 is primarily a cytosolic GEF, where it activates Cdc42 (Tay *et al*. 2018). Rather our data suggest that Cdc15 recruits Gef1 to the cortical patches to promote bipolar growth. During interphase Cdc15 is localized to the endocytic patches where it promotes vesicle internalization (Arasada and Pollard 2011). In the hypermorphic mutants, cdc15-27A-GFP levels are elevated at cortical patches (Roberts-Galbraith *et al*. 2010). Correspondingly, these mutants also show Gef1 localization to these patches. Moreover, Gef1 localization at the cortex is quite prominent in these mutants. In agreement with increased Gef1 cortical localization, we also observe increased Cdc42 activation at both the cell poles resulting in increased bipolarity. A recent paper demonstrates that Gef1 regulates Cdc15 by controlling the size and lifetime of Cdc15 cortical patches (Onwubiko *et al*. 2019). Above, we present data that demonstrate Cdc15 is upstream of Gef1. These two observations are not contradictory, but rather reveal an elegant regulatory pattern: Cdc15 recruits Gef1 to endocytic patches, where Gef1 in turn regulates the size of the Cdc15 patch via Cdc42 activation. Our observation that Gef1-tdTomato and Cdc15-27A-GFP do not perfectly colocalize at the cortex can be explained by the following model. Cdc15 initially recruits Gef1 to endocytic patches at the cortex, resulting in colocalization. Once Gef1 facilitates patch internalization, Cdc15 is lost from the cortex while Gef1 still persists. Further investigations will determine how Gef1 mediated Cdc42 activity regulates Cdc15 cortical patch lifetime. Given the abundance of Gef1 in the cytoplasm, small levels of Gef1 are not easily detectable at cortical patches. Gef1 localization to the cortical patches and the cortex may be enhanced by the increased abundance of cdc15-27A at cortical patches.

Since we established that *cdc15* promotes Gef1 localization to the division site, and that *cdc15-27A* enhances Gef1 localization to cell tips, we asked whether *cdc15-27A* would similarly result in precocious Gef1 localization to the division site. Indeed, we find that Gef1 localizes to node like patches at the cell equator in interphase cells. In keeping with this observation, Gef1 was detected during early stages of cdc15-27A-labelled ring assembly, while it is not yet detectable at the comparable time in *cdc15*^+^ cells. Further, we show that precocious Gef1 localization results in premature Cdc42 activation. Finally, we show that the *cdc15* mutant phenotypes, which arise from the mis-regulation of Cdc42, are *gef1*-dependent. While *cdc15-27A* cells are mainly bipolar, loss of *gef1* in these cells reverts to the *gef1Δ* monopolarity phenotype. Cdc42 activation at the division site is also delayed in *cdc15-27A gef1Δ*. Together these results indicate that *gef1* is epistatic to *cdc15*.

The mechanistic understanding of factors that control Gef1 localization is sorely lacking. Aside from the observation that the N-terminus of Gef1 is required for its localization to the membrane, no other factors have been identified (Das *et al*. 2015). It has also been reported that Gef1 activates Cdc42 with the help of N-BAR Hob3 protein interaction (Coll *et al*. 2007). Gef1 is a homolog of the mammalian GEF TUBA and contains an N-BAR domain (Das *et al*. 2015). However, previous reports show that the Gef1-N-BAR domain is not required for its localization to the division site, nor is Hob3 required for Gef1 localization (Supplementary Fig. S2) (Das *et al*. 2015). In contrast, the mechanism removing Gef1 from the membrane has been elucidated. Gef1 is phosphorylated by Orb6, generating a 14-3-3 binding site that results in Gef1 removal by Rad24 (Das *et al*. 2009; Das *et al*. 2015). Here, we identify Cdc15 as a factor that promotes Gef1 localization to both the cell tips and division site. A recent study indicates that Gef1-mediated Cdc42 activation regulates endocytosis, by controlling the lifetime of Cdc15 on endocytic patches (Onwubiko *et al*. 2019). The role of endocytosis in cell polarity is well established and its role in cytokinesis is increasingly recognized (Wang *et al*. 2016; Onwubiko *et al*. 2019). Taken together, this suggests that Cdc15 promotes Gef1-mediated Cdc42 activation to regulate endocytosis to promote both bipolar growth and cytokinesis. We find that Cdc15 promotes the localization of Gef1, but not Scd1. These studies begin to explain how, through differential regulation and localization, two GEFs of the same GTPase can exhibit distinct phenotypes.

## ACKNOWLEDGEMENTS

We thank Kathleen Gould and Fred Chang for generously providing strains. This work was funded by National Science Foundation, grant #1616495.

---

**Supplemental Figure 1.**
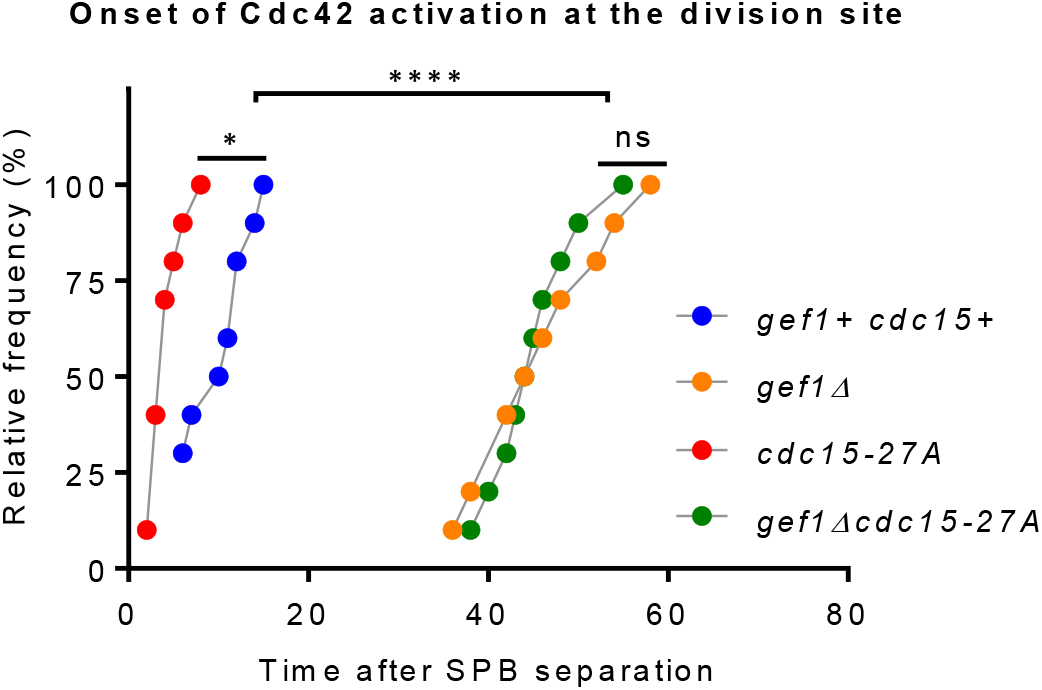
Precocious Cdc42 activation at the division site in *cdc15-27A* cells is *gef1*-dependent. Quantification of Cdc42 activation at the division site from time lapse images the indicated genotypes expressing CRIB-3xGFP and Sad1-mCherry. ****, p<0.001, *, p<0.05, ns=not significant, one-way ANOVA with Tukey’s multiple comparisons post hoc test.

**Supplemental Figure 2.**
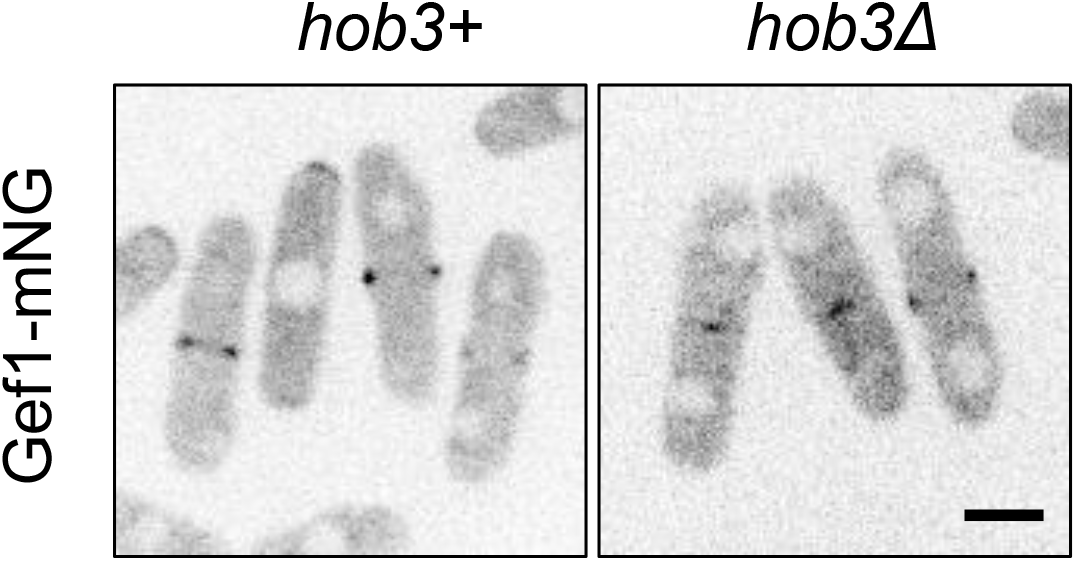
Gef1 localization is not impaired by loss of *hob3*+. Inverted medial plane images showing Gef1-mNG localization in the indicated genotypes. Scale bar represents 5μm.

**Table S1.**
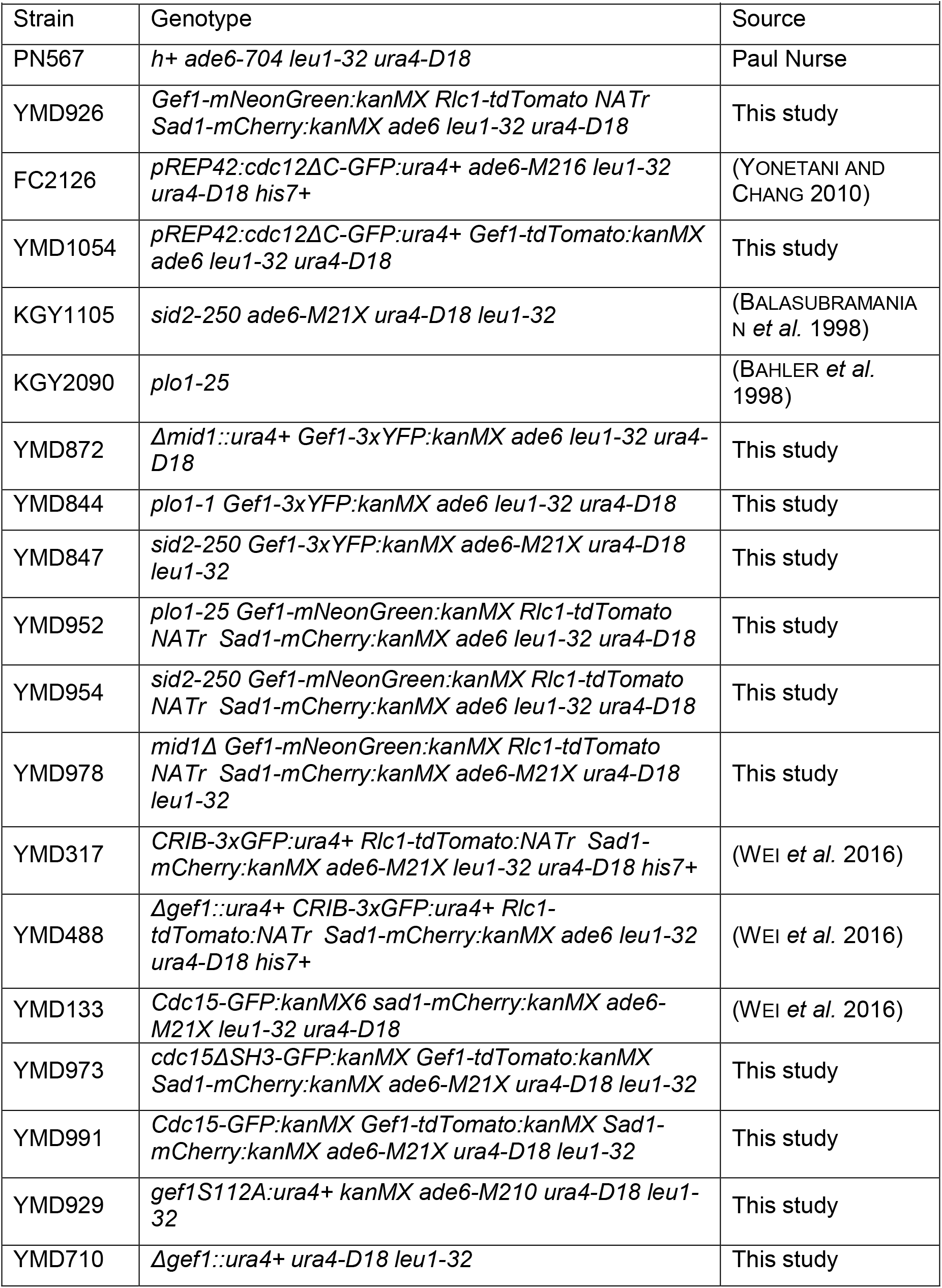
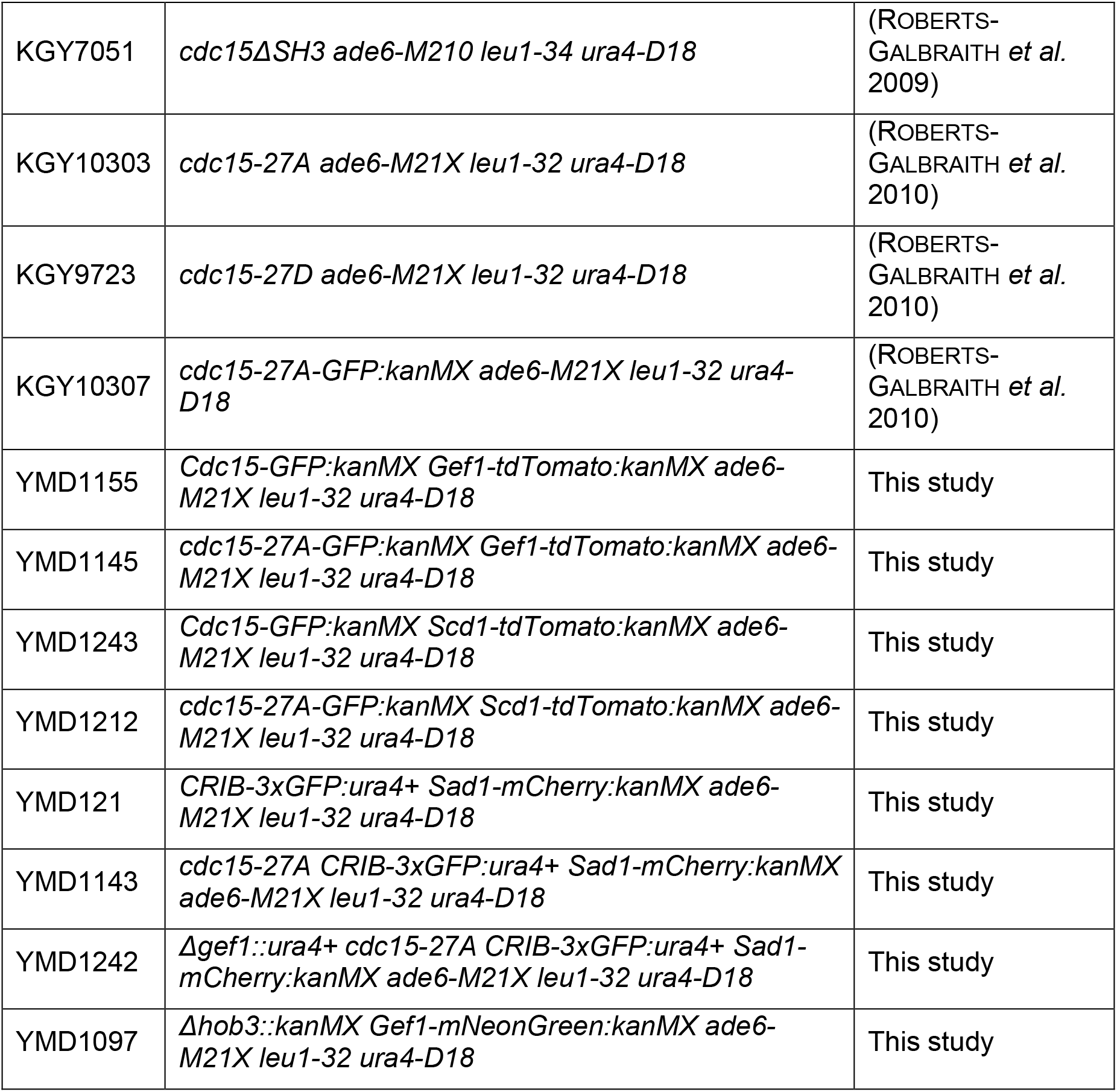
Strains list.

